# Evidence for independent retroviral syncytin-like Env endogenization in non-placental chondrichthyans

**DOI:** 10.64898/2026.05.06.723177

**Authors:** Ella Proudley, Ian G Reddin, Jane K Cleal, Rohan M Lewis, Davis Laundon

## Abstract

Viviparity and placentation are remarkable examples of convergent evolution across vertebrates. The evolution of the uniquely intimate mammalian placenta has been associated with the repeated independent capture of fusogenic retroviral Env proteins, called syncytins. Research into syncytin capture has therefore been predominantly focused on resolving their central role in mammalian placentation. As such, the presence of syncytin-like Env proteins outside of mammals, and their role in non-placental physiological contexts, remain much less understood. We expanded this understanding by systematically surveying genomes from 36 chondrichthyan species (sharks, rays, skates, and chimaeras), which display a wide range of independently evolved placental and non-placental reproductive strategies, for the presence of syncytin-like Env genes. We identified 295 candidate syncytin-like Env proteins from 16 chondrichthyan species, with a subset displaying conserved fusogenic domains, structural homology with known syncytins, and genomic signatures of endogenization. Using transcriptomic data from the model catshark *Scyliorhinus canicula*, we found that syncytin-like Env genes are transcriptionally active in diverse adult tissue types. Using two closely related species of *Squalus* (spiny dogfish), we present evidence that endogenized Env genes are syntenically conserved, indicative of vertical transmission from a common ancestor before species divergence. Notably, we detected no candidates in any placental shark genome, suggesting that syncytin-like Env capture is not a feature of shark placentation. Our findings expand the known phylogenetic breadth and functional scope of syncytin-like Env protein endogenization beyond mammalian placentation, providing a solid foundation for future investigations into the wider role of retroviral capture in vertebrate biology and evolution.

## Introduction

Viviparity (live birth) has evolved at least 150 separate times in vertebrates [1, 2], making it one of the most remarkable cases of convergent evolution in animal reproduction. Switching parity mode from oviparity (egg laying) to viviparity is often coupled to innovations in nutrient provisioning, moving from lecithotrophy (nutrient provisioning from a deposited, finite yolk source) to matrotrophy (continuous nutrient provisioning from the mother to the embryo across gestation) [3-5]. The most complex form of matrotrophy is placentotrophy, whereby maternal nutrients are provisioned to the embryo via an embryonic placenta [6, 7]. Like viviparity, complex nutritive placentas have emerged multiple times independently in vertebrates, often from diverse precursory tissues [8]. Mammals are notable among vertebrate groups as every species (except for monotremes) displays placentotrophic viviparity, which originated only once in the common ancestor of all therian mammals [9-11]. Many mammals, including humans, exhibit a uniquely intimate ‘hemochorial’ form of placental interface, whereby the placenta is in direct contact with maternal blood [12, 13]. Ancestral trait reconstruction has repeatedly shown that the common ancestral mammalian placenta was hemochorial [10, 14-16].

Key to the evolution of the intimate hemochorial placental interface in mammals is the co-option of retroviral envelope (Env) genes called ‘syncytins’ [17-20]. In retroviral genomes, Env glycoproteins are responsible for membrane anchorage and fusion between the host cell and retroviral envelope [21]. However, if the host germline is infected with proviruses, these sequences can be integrated into the host genome, fixed, and vertically transmitted to offspring in a process called ‘endogenization’ [17, 22]. Endogenized Env sequences have been co-opted into mammalian placentation, where their fusogenic properties have been exapted for cell-cell fusion in the placental interface [17]. Mammalian syncytins, such as human syncytin-1 and -2 or the independently captured mouse syncytin-A and -B, mediate the fusion of mononucleated cytotrophoblast cells into a syncytial trophoblast interface [23-25] (Figure 1A). The syncytial trophoblast interface is thought to facilitate intimate hemochorial pregnancies by reducing the risk of vertical pathogen transmission [26, 27] and suppressing rejection by the maternal immune system [28, 29].

**Figure 1.**
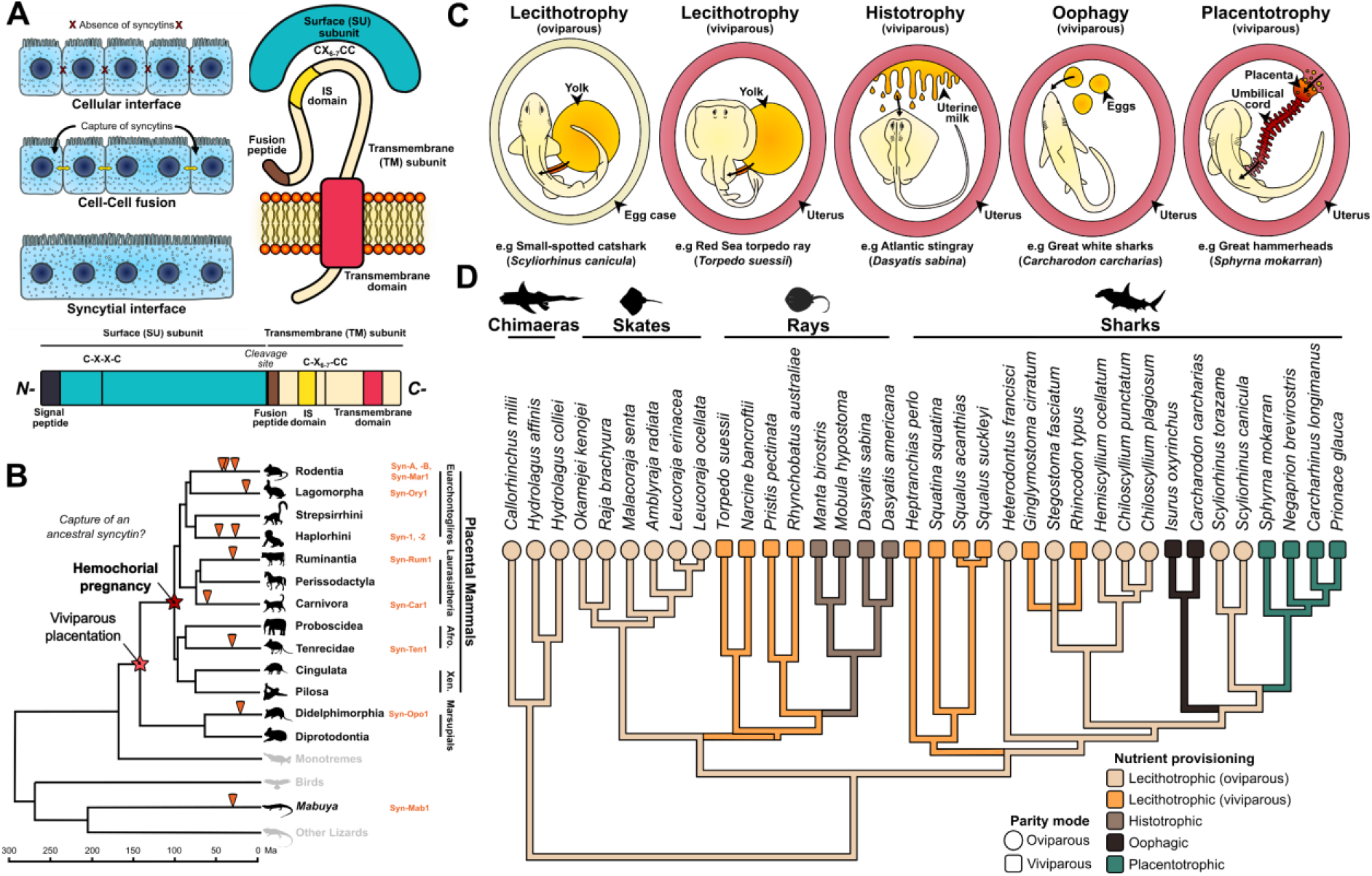
Chondrichthyans present a powerful natural system to investigate the independent capture and evolution of retroviral Env proteins outside of the mammalian placenta. (A) When endogenized syncytins are expressed in the mammalian placenta, their fusogenic activity promotes the syncytialisation of the trophoblast interface (left). Shown is a schematic of the canonical Env protein structure (right) and peptide sequence (bottom). Schematic and peptide sequence adapted from [30]. (B) Syncytin proteins have been independently endogenized at least ten separate times in mammals, and once in *Mabuya* lizards (adapted from [30]). (C) Diagrammatic summary of chondrichthyan parity modes and maternal nutrient provisioning strategies with example species for which genomes are available. (D) Phylogenetic distribution and nutrient provisioning strategies of the 36 chondrichthyan species whose genomes we surveyed for syncytin-like Env sequences. Phylogeny based on [37] and trait data taken from [34].

The origin of uniquely intimate mammalian placentation may have been driven by the capture of an ancestral syncytin [17, 19]. Such is the central role of syncytin capture in mammalian placenta diversification, that syncytins have been captured at least 10 times independently across the mammals (and once in the placental lizard *Mabuya*) (Figure 1B), each time conferring fusogenic and more intimate attributes to placentation [30]. Syncytin-like endogenized Env sequences have also been detected in bony fish, where they are known as percomORFs [31, 32]. However, the strong research emphasis on syncytin capture and the evolution of mammalian placentation has limited our understanding of the roles played by independently captured syncytin-like Env proteins in less intimate placentation, as well as in non-placental and non-mammalian reproductive systems. As a result, our knowledge of the presence of syncytin-like Env proteins outside of the mammals, and their association with non-placental nutrient provisioning strategies, remains limited. Addressing this knowledge gap is necessary if the unique innovations of mammalian placentation are to be placed in their proper context, and the role of retroviral Env capture is to be understood in the convergent evolution of vertebrate reproductive biology more broadly.

Chondrichthyans (sharks, skates, rays, and chimaeras) present a powerful natural system to investigate the evolution of vertebrate reproductive strategy [33, 34]. Chondrichthyans are a large and ancient (∼420 Ma) clade where viviparity has evolved at least 12 separate times from oviparous ancestors and all major modes of matrotrophic nutrient provisioning are present from six independent origins [8, 34] (Figure 1C). Within these modes of matrotrophy is a single origin of placentation in the Carcharinidae (requiem sharks) and their viviparous kin within the Carcharhiniformes (ground sharks) [34]. Unlike the ancestral mammalian placenta, shark placentas (derived from the yolk sac) are not in direct contact with maternal blood but simply apposed to the uterine epithelium (they are epitheliochorial, not hemochorial) [8, 33]. However, the presence of syncytin-like Env sequences in chondrichthyans, and their association with shark placentation and alternate modes of nutrient provisioning, is not well understood.

Although chondrichthyan genome sequencing has been less extensive than that of mammals, we now have genomic information for species across the entire clade, spanning species with diverse parity modes and nutrient provisioning strategies [35, 36] (Figure 1D). These publicly available genomic data present a valuable resource to investigate the capture of syncytin-like Env proteins in non-mammalian taxa and elucidate their roles outside of intimate placentation. Here, we systematically screen the genomes of 36 chondrichthyan species against known syncytin and syncytin-like proteins to identify novel candidates for independently endogenized retroviral Env sequences. Using a combination of sequence analysis, 3D protein structure prediction, and publicly available transcriptomic datasets, we go on to demonstrate evidence for independent syncytin-like Env endogenization, expression, and vertical transmission in non-placental sharks.

## Methods

### Chondrichthyan phylogeny

The phylogenetic tree of the chondrichthyan species for which there are genomes analysed in this study (Figure 1D) was built based on the phylogeny outlined by [37] using the online resource at: https://vertlife.org/data/sharks/. Consensus trees were compiled from 1,000 tree replicates using the *consensus*.*edges* function in the R package *Phytools* v.2.1.1 [38] run with R v.4.4.0 implemented in RStudio v.2024.4.1.748. Trait data for parity and nutrient provisioning for each species was taken from [34].

### Identification of syncytin-like sequences in chondrichthyan genomes

To identify novel syncytin-like Env genes in chondrichthyans, we first compiled a database of all sequenced genomes from GenBank assigned to the Chondrichthyes clade using the NCBI Taxonomy Browser. This resulted in 63 genomes (38 of which were reference genomes) from 36 species of chondrichthyans (Figure 2). We then compiled a database of query sequences (Supplementary File 1) from ten groups of syncytins independently captured in mammals and the *Mabuya* lizards known to play a fusogenic role in the placenta (Figure 1B), in addition to syncytin-like Env genes identified in hyenas (Hyena-Env2) [30], bony fish (percomORF’s) [31], and monotreme mammals (env-Tac’s) [39].

**Figure 2.**
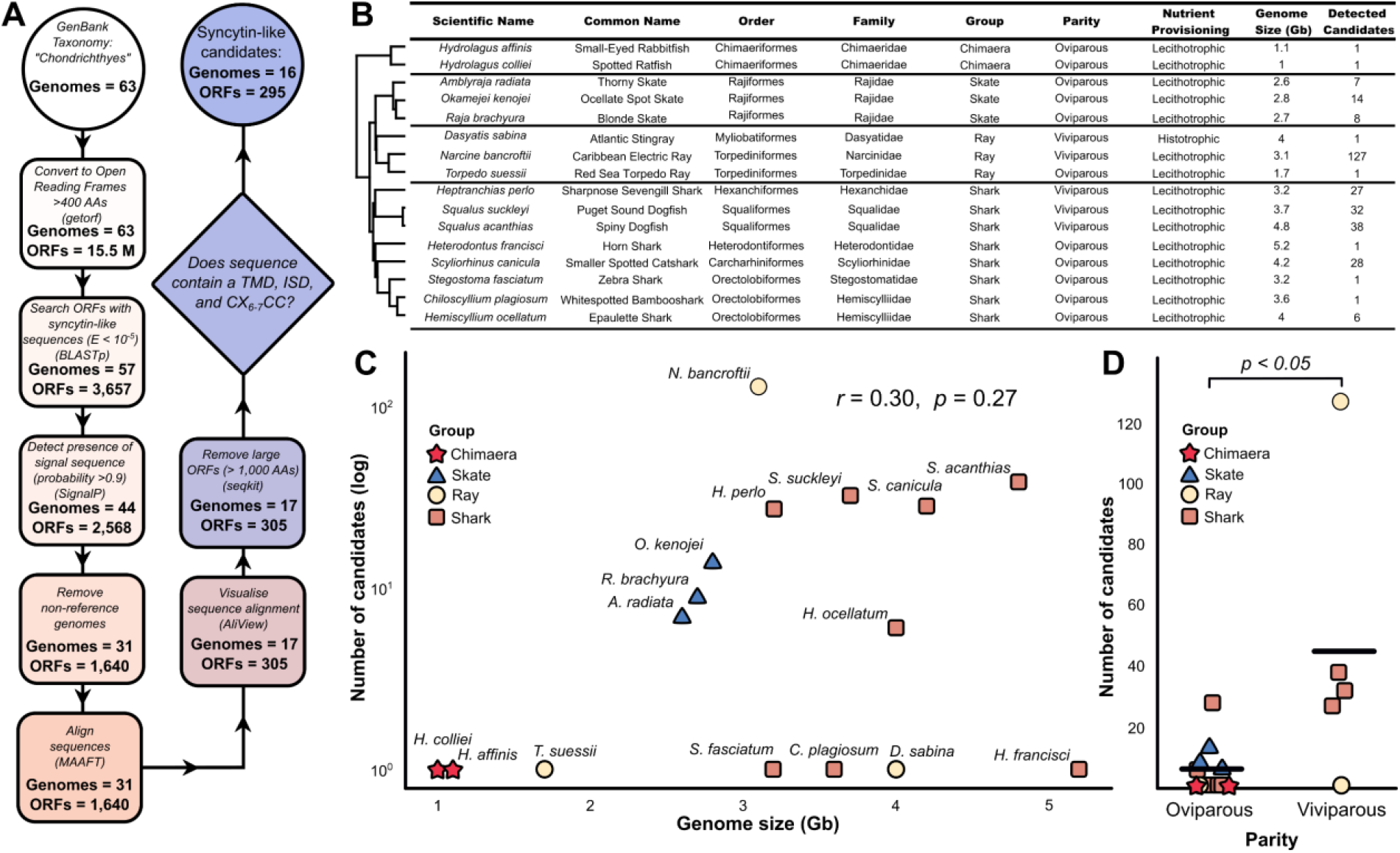
A systematic search of known syncytins detected novel candidate Env proteins in chondrichthyans. (A) Flow chart summarising the methodology and results at each stage of our bioinformatic pipeline to search known syncytins against chondrichthyan genomes. (B) Our systematic search identified candidate syncytin-like Env proteins in 16 chondrichthyan species. (C-D) There was no correlation between the number of identified candidates with genome size (C), however we detected more candidates in viviparous species than oviparous species (D).

The chondrichthyan genomes were converted into open reading frames (ORFs) using the *getorf* function (EMBOSS 6.6.0.0) [40] with a minimum nucleotide threshold of 1200 bases (∼400 amino acids) based on the lengths of previously known syncytin proteins, which were output as translated amino acid sequences. The compiled syncytin query sequences were searched against the chondrichthyan genome ORFs using BLASTp [41] with a minimum E-value threshold of 10^-5^. Positive ORF hits were then screened for the presence of a functional signal peptide at the N terminus using SignalP v.6.0 [42], and sequences below a threshold of 0.9 (90% likelihood) were excluded from further analysis.

Positive sequences were aligned using MAAFT 7.490 [43] (Figure 3A&B). Some ORFs were much larger than expected syncytins, so at this stage, the EMBOSS *seqkit* function was used to remove sequences >1,000 amino acids. Following this, the remaining aligned sequences were trimmed using the EMBOSS *gappyout* function. The sequence alignment was viewed in AliView 1.28 [44] and manually inspected for the presence of conserved domains necessary for Env protein function. These included the R-X-X-R putative furin cleavage site delimitating the surface (SU) and transmembrane (TM) subunits, the immunosuppressive domain (ISD), the conserved C-X_6-7_-C (CC) motif, and the hydrophobic TM subunit itself. Any sequence meeting all these criteria was considered a candidate syncytin-like Env protein.

**Figure 3.**
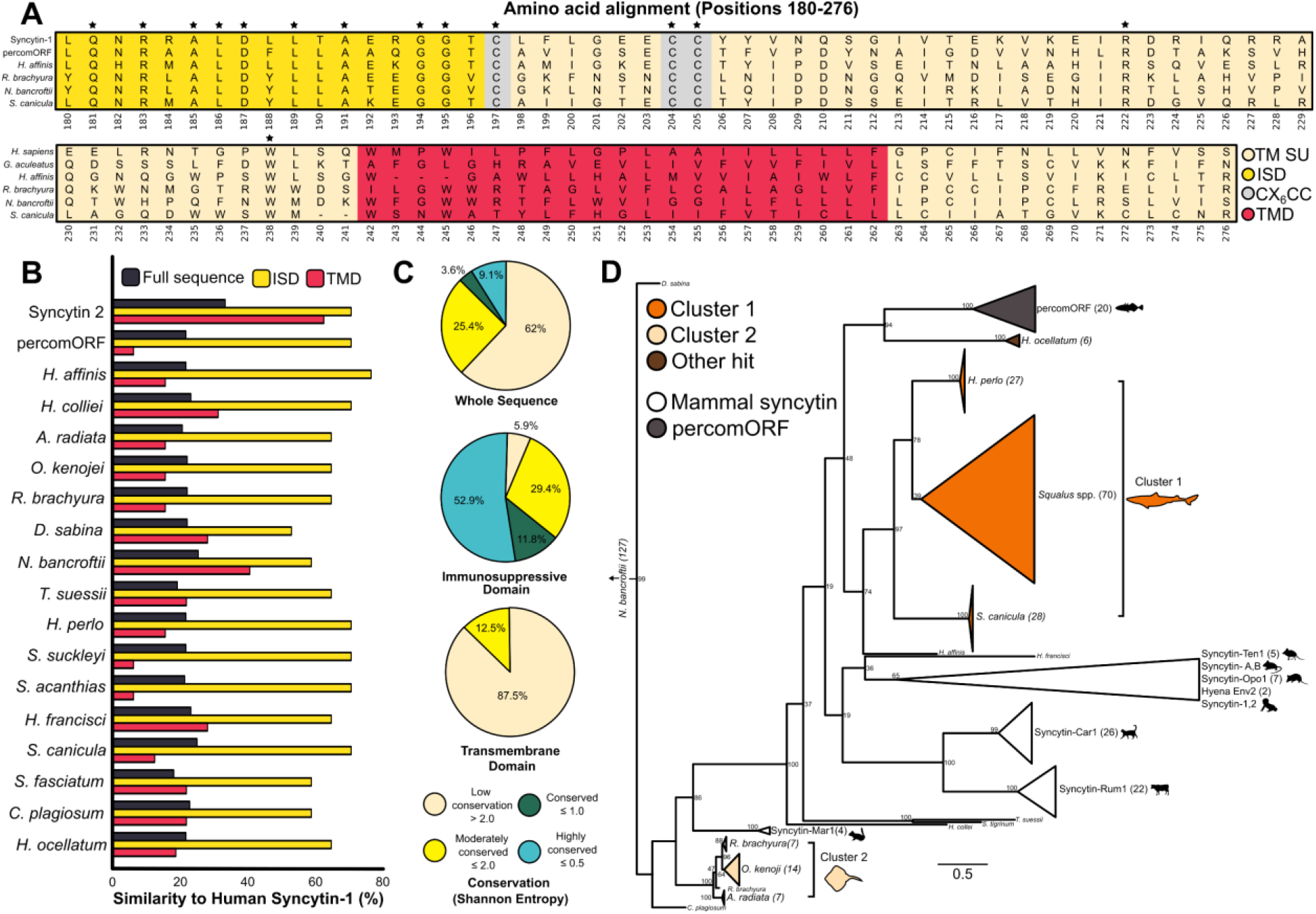
Identified candidate syncytin-like Env proteins shared a highly conserved immunosuppressive domain and formed two distinct phylogenetic clusters. (A-C) Multiple sequence alignment of the transmembrane subunit (TM SU) consensus sequences of syncytin-like Env protein candidates from four chondrichthyan species with human syncytin-1 and stickleback percomORF shows a highly conserved immunosuppressive domain (ISD). Asterisks in (A) show sites where amino acids are identical across all species. (B-C) The ISD in chondrichthyan candidates shows much higher sequence similarity with human syncytin-1 than either the full sequence or the TMD (B), and Shannon Entropy analyses shows that the ISD is much more highly conserved (C). (D) Maximum likelihood (ML) phylogeny of chondrichthyan candidates showed two distinct clusters. Cluster 1 was composed of shark species and grouped with percomORF sequences next to mammalian syncytins. Cluster 2 was composed of ray species and clustered as an outgroup to the other sequences. Candidates from other species did not cluster and were found scattered throughout the tree. Branch length represents the average number of substitutions per site (scale bar). Tree run with 1,000 bootstraps.

### Statistical analysis

Prior to analysis, data was assessed with a Shapiro and Levene’s test for homogeneity and normality respectively, and an appropriate statistical test selected. The relationship between genome size and syncytin-like Env candidate number was assessed using a Pearson’s correlation (Figure 2C). Comparison of candidate number by parity mode was assessed using a Mann-Whitney-U test (Figure 2D). All statistical comparisons were conducted using the Python package *scipy* v1.11.4 [45] run with Python 3.11.7 implemented in Jupyter Notebook 7.0.8.

### Sequence analysis

Amino acid conservation was quantified using Shannon entropy calculations, where low entropy (≈ 0) indicates high conservation at the position, high entropy (≤ 4.32) indicates high variability (Figure 3C). RAxML-NG 8.2.12 [46] was used to produce a maximum-likelihood tree of candidate chondrichthyan Env proteins and known syncytins run with 1,000 bootstraps (Figure 3D), which was visualised in FigTree 1.4.3. Hydrophobicity was calculated using the Kyte-Doolittle hydrophobicity scale [47] (Figure 4A&B). All amino acid sequence analyses were conducted and visualised using *Biopython, NumPy, matplotlib, pandas*, and *seaborn* packages run with Python 3.11.7 implemented in Jupyter Notebook 7.0.8. 3D protein architecture of candidate syncytin-like Env genes were predicted using AlphaFold2 [48], implemented and visualised using UCSF ChimeraX 1.8 [49] (Figure 4C-E). Amino acid conservation, hydrophobicity, and structural analyses were conducted on consensus sequences of each species candidate copies, generated using the EMBOSS *cons* function.

**Figure 4.**
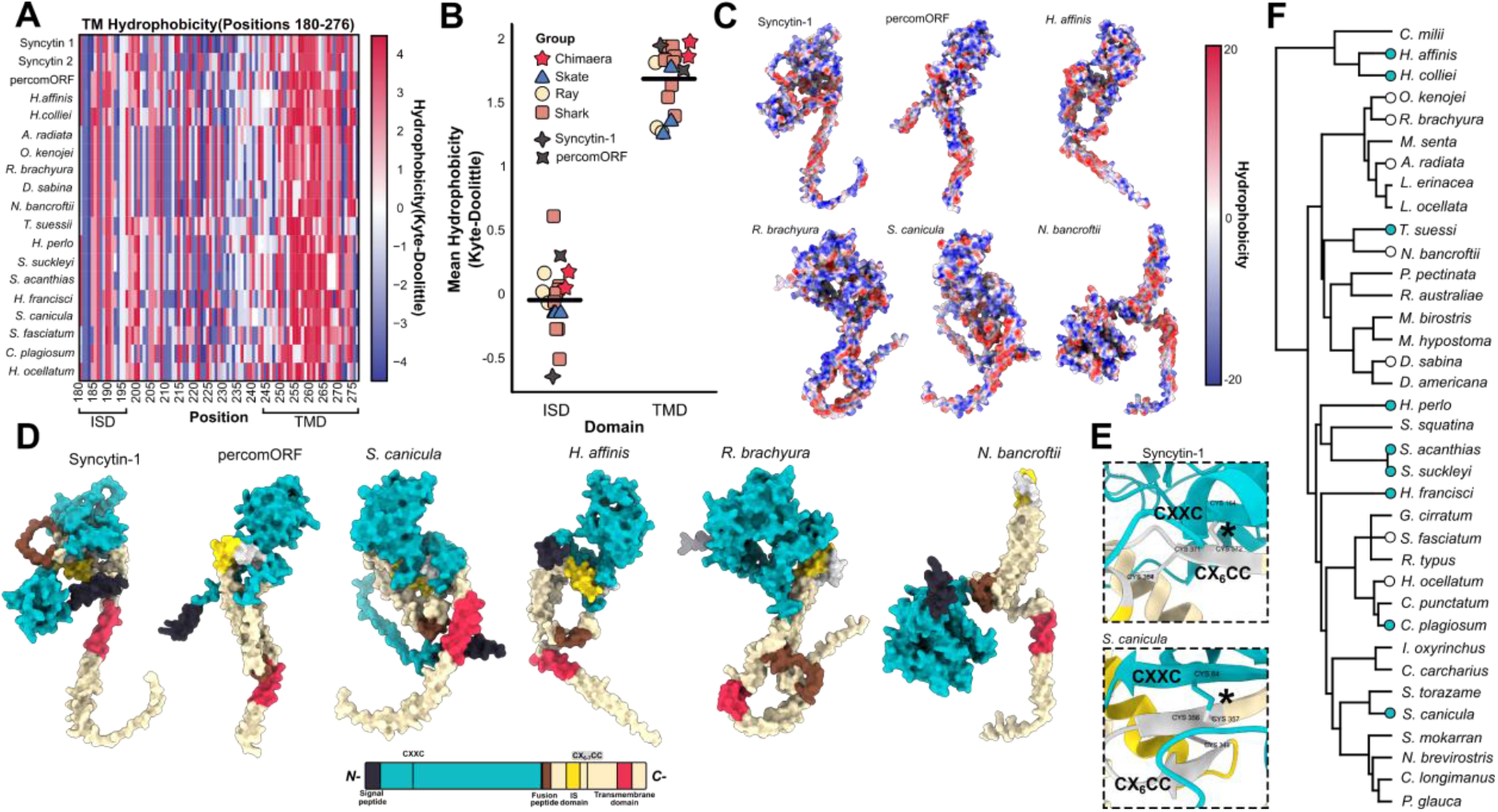
3D structure prediction of syncytin-like Env sequences demonstrates the presence of a hydrophobic transmembrane domain and key disulfide bond necessary for fusogenic function. (A-C) The TMD in candidates is much more hydrophobic than the ISD (A-B) and hydrophobic residues localise to the site of membrane anchorage in example candidates (C), as would be necessary for fusogenic function. (D-F) 3D structure prediction of some candidate proteins (cyan circles in F) showed conformational conservation (D) and a canonical disulfide bond (E) as seen in mammalian syncytins and percomORFs. (F) Candidates which did not meet these criteria were excluded from further analysis (white circles).

### Env expression in RNA-seq datasets and Gag-Pol analysis

A total of 245 RNA-Seq datasets were available from diverse *S. canicula* tissue types (Figure 5A&B). Candidate genomic coordinates were harmonized to match reference genome FASTA contig identifiers. Where necessary, GenBank contig accessions associated with candidate loci were translated to corresponding RefSeq contig accessions using the NCBI assembly sequence report (for *S. canicula* genome sScyCan1.1). Harmonized coordinates were exported in BED6 format, with strand determined from interval orientation. Candidate nucleotide sequences were extracted directly from the reference genome FASTA using strand-aware extraction, yielding an intronless candidate FASTA suitable for transcript-level quantification. Expression of candidate loci was quantified from adult RNA-seq data using Salmon (v1.11.4) [50]. A targeted indexing strategy was employed in which candidate sequences were quantified either alone or using a decoy-aware index that included the full reference genome as decoy sequences to reduce spurious alignments from repetitive regions. Salmon selective alignment was used with automatic library type detection, and sequence- and GC-bias corrections were enabled.

**Figure 5.**
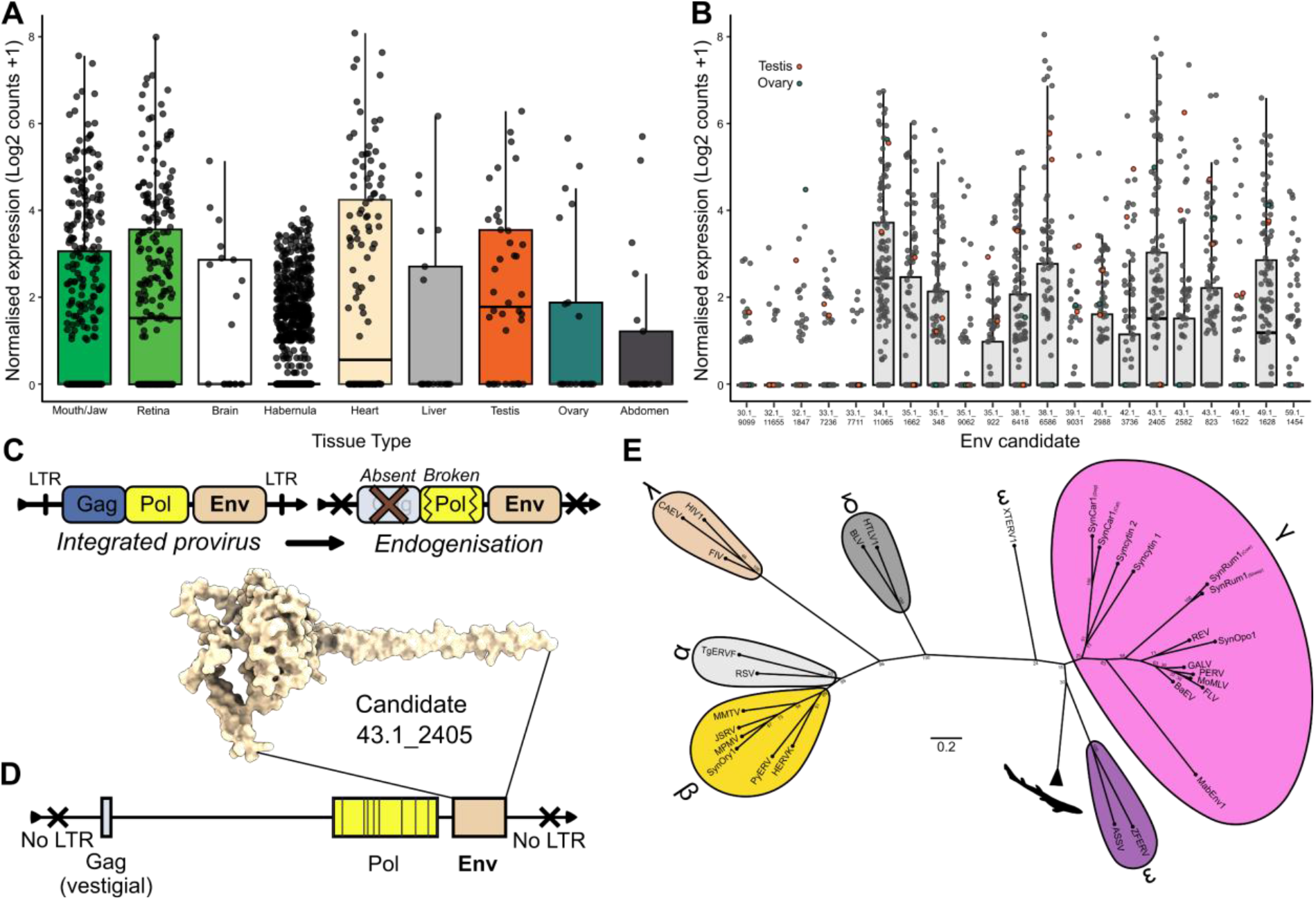
Syncytin-like Env genes in the model catshark *Scyliorhinus canicula* are expressed in diverse tissue types and show genomic signatures of endogenization. Env candidates identified in the *S. canicula* genome were detected as expressed in available RNA-seq dataset from diverse adult tissue types (A). Candidates in which expression was detected (B) were carried forward for further analysis. (C) Diagrammatic summary of key elements of domain change during retroviral endogenization. The integrated provirus maintains Env function while the Pol gene is disrupted by stop codons, the Gag becomes vestigial or absent, and the LTRs disappear. LTR = long tandem repeats. (D) Example candidate from *S. canicula* (43.1_2405) showing genomic signatures associated with Env endogenization. Vertical lines in the Pol gene represent stop codons. (E) Unrooted ML tree of Pol genes associated with endogenous syncytins and known retroviruses against candidates indentified in *S. canicula* (shark silhouette), suggesting an origin from an ancestral Epsilonretrovirus. Branch length represents the average number of substitutions per site (scale bar). Tree run with 1,000 bootstraps.

Candidate retrotransposon loci are expected to be lowly expressed and may be subject to multi-mapping ambiguity. Therefore, expression detection was evaluated using conservative but permissive thresholds. Per sample, candidates supported by ≤ 3 estimated reads were considered not detected. Candidates supported by 5–10 estimated reads were classified as showing weak evidence of transcription. Candidates supported by ≥ 10 estimated reads in each sample were considered expressed, and those supported by ≥ 50 estimated reads were considered robustly expressed. For analyses requiring quantitative stability, only candidates with ≥ 10 estimated reads in two or more adult samples were retained. Expression estimates were interpreted as evidence of transcription consistent with the candidate sequence, with appropriate caution given the repetitive nature of retrotransposon-derived loci.

To evaluate nearby retroelement-derived coding signals, such as the presence of Gag and Pol genes (Figure 5C&D), genomic windows (± 25 kb) around each anchor locus were translated by six-frame stop-to-stop ORF scanning, and translated ORFs above the minimum length threshold were screened with HMMER [51] *hmmscan* against a focused retroelement panel including PF03732 (Gag-like), PF00077 (retroviral protease-like), PF00078 (reverse transcriptase), PF00075 (RNase H), and PF00665 (integrase core/rve-like). Because PF00665/rve-like matches are not uniquely diagnostic of retroviral Pol elements, loci were interpreted using a strict architecture scheme based on the combination of nearby domain types. Candidate loci were classified as *gag_and_rt_supported_pol, multi_domain_pol, rt_supported_pol_only*, or *integrase_like_only*. To test whether candidates retained recoverable LTR (long tandem repeats) retrotransposon structures, larger windows spanning ± 50 kb around each anchor locus were screened using an LTR-detection workflow based on GenomeTools v.1.6.6 *ltrharvest*, followed by filtering with *LTR_retriever* v3.0.5. Candidate-level summaries recorded whether an intact LTR retrotransposon candidate was present within the window and whether the anchor ORF fell within such a structure. Retrieved Pol sequences in *S. canicula* were clustered in a maximum likelihood tree as in Figure 3D above with a dataset of known retroviral and endogenous Pol sequences [30] (Figure 5E).

### Pairwise synteny and ORF projection analysis

Candidate ORFs from *Squalus acanthias, Squalus suckleyi*, and *S. canicula* were compiled as anchor loci for local retroelement-context analysis. For *S. canicula*, chromosome accessions were remapped from GenBank to RefSeq identifiers before downstream analysis. In the current strict-domain summary, *n* = 38 loci were analyzed for *S. acanthias, n* = 21 for *S. canicula*, and *n* = 31 for *S. suckleyi*. Comparative synteny analysis was performed using the genome assemblies for *S. canicula* (sScyCan1.1), *S. acanthias* (ASM3039002v1), and *S. suckleyi* (GSC_Ssuck_1.0). Directly comparable gene annotations were not available across assemblies, so a whole-genome alignment strategy was used rather than a gene-anchor collinearity pipeline. Pairwise genome alignments were generated with minimap2 [52] in assembly-alignment mode and retained in PAF (pairwise mapping format) format with CIGAR information. Initial comparisons included *S. canicula* vs *S. acanthias*; *S. canicula* vs *S. suckleyi*i; and *S. acanthias* vs *S. suckleyi*. The intergeneric comparisons yielded sparse short matches and were not suitable for robust genome-scale interpretation, whereas the *S. acanthias–S. suckleyi* comparison produced extensive high-confidence alignments and was therefore selected for downstream locus-level analysis.

For the focal *S. acanthias–S. suckleyi* comparison, whole-genome alignments were visualised as ordered PAF dot plots, retaining long, high-confidence alignment blocks of ≥ 100 kb with mapping quality ≥ 20. *S. suckleyi* scaffolds were assigned to their dominant *S. acanthias* chromosome based on total aligned base pairs and ordered by chromosome assignment and aligned position to generate an interpretable synteny map (Figure 6). *S. suckleyi* candidate ORF intervals were then projected into *S. acanthias* coordinates using overlapping primary PAF alignment blocks. For each ORF, the best overlapping alignment block was chosen based on overlap with the ORF interval, followed by mapping quality and alignment length, and coordinates were projected into the *S. acanthias* space. Projected intervals were compared with the observed *S. acanthias* ORF coordinates and classified as overlap, nearby, *same_chr_only*, or unresolved. Because projection was based on whole-genome alignment blocks rather than base-level gap-aware liftover, overlap calls were interpreted as evidence of shared syntenic location rather than identical nucleotide-level insertion sites. Local conservation was further assessed using ORF-level coverage and flanking-window (± 10 kb) alignment summaries for each projected locus.

**Figure 6.**
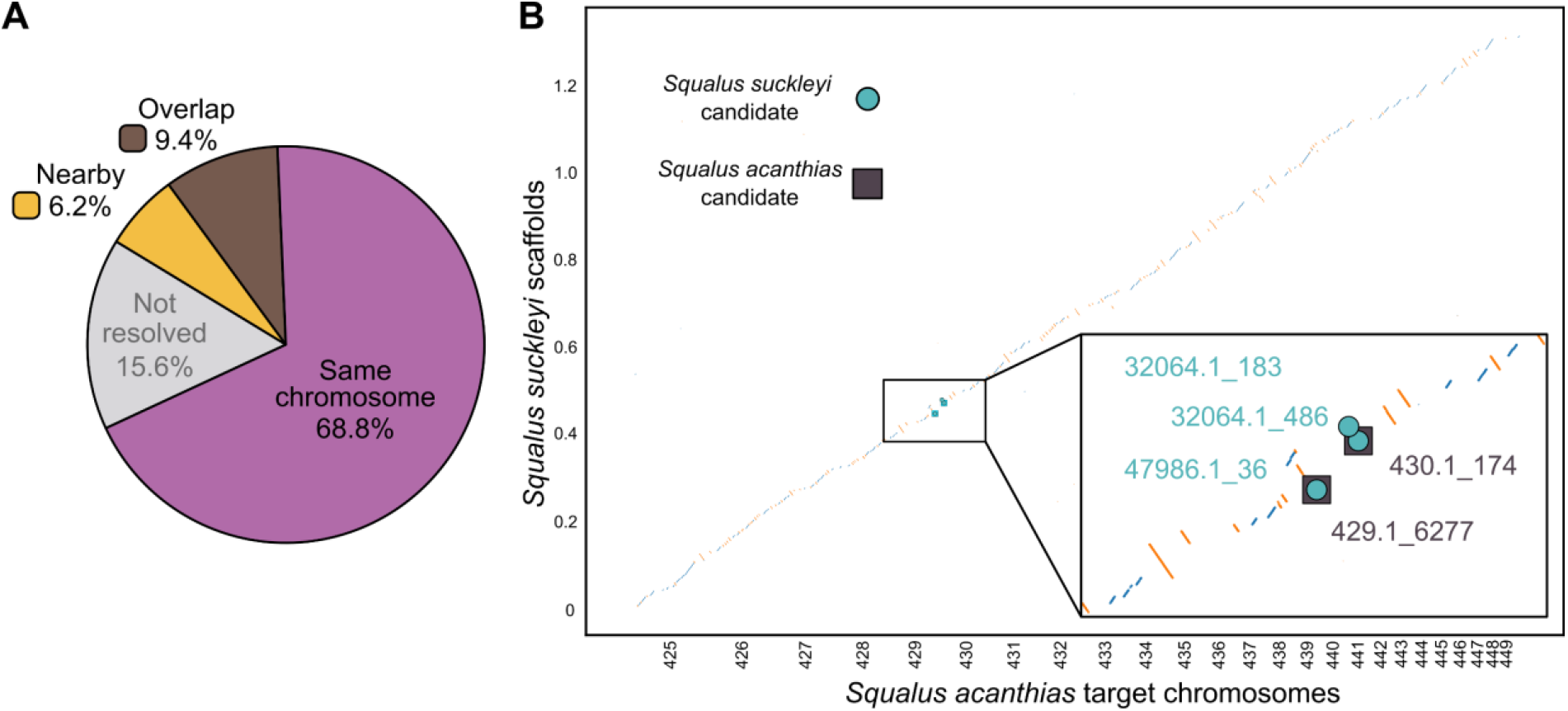
Endogenized syncytin-like Env genes show syntenic conservation in two *Squalus* species indicative of vertical transmission from a common ancestor. (A) Percentage distribution of syncytin-like Env candidates identified in *S. suckleyi* based on their syntenic conservation with *S. acanthias* candidates from whole genome alignment. (B) Illustrative co-localisation of overlapping and nearby syncytin-like Env candidates (inset) overlaid on the *S. acanthias*-*S. suckleyi* aligned genome space.

## Results

Our pipeline (Figure 2A) identified a total of 295 syncytin-like Env ORF’s in 16 species of chondrichthyans (8 sharks, 3 rays, 3 skates, and 2 chimaeras), none of which were placental (Figure 2B; Supplementary File 2). Of the species with candidate syncytin-like Env proteins, 11 were oviparous (all with lecithotrophic nutrient provisioning) and 5 were viviparous (4 with lecithotrophic provisioning, and 1 with histotrophic nutrient provisioning) (Figure 2B). We did not detect any candidates in chondrichthyans with oophagous or placental nutrient provisioning. In species with candidate sequences, there was no correlation between candidate number and genome size (*r* = 0.30, *p* = 0.27) (Fig. 2C), however viviparous species did display a higher candidate number than oviparous species (*p* < 0.05) (Figure 2D).

All candidate TM units displayed a canonical immunosuppressive domain, transmembrane domain, and conserved C-X_6_-CC cysteine-containing region (essential for the formation of disulfide bonds with the SU C-X-X-C motif, linking the SU and TM units) (Figure 3A). Overall, identified candidates displayed a 22.6 ± 3.2% (mean ± SD) amino acid similarity to human syncytin-1 (for reference, human syncytin-2 is 33.3% similar to syncytin-1), with the immunosuppressive domain being much more similar (66.0 ± 5.9%) than the transmembrane domain (21.4 ± 13.7%) (Figure 3B&C). Chondrichthyan candidates displayed two discrete clusters when analysed by maximum likelihood phylogeny (Figure 3D). The first cluster is represented by sequences in the shark species *H. perlo, S. acanthias* and *S. suckleyi*, and *S. canicula*. The second cluster is represented by sequences in the skate species *O. kenojei, R. brachyura*, and *A. radiata*. The first cluster of shark sequences clustered more closely to known mammalian syncytin and *percomORF* sequences than it did to the second. The large number of sequences in *N. bancroftii* (127) clustered heterogeneously outside all other sequences, with poor bootstrap support, and as such were excluded from the tree in Figure 3D.

Although the amino acids in the TM domain were less conserved than in the ISD, the residues were still more hydrophobic overall as would be expected for the functional membrane anchorage of an envelope protein (Figure 4A&B). Visualisation of hydrophobicity against predicted protein structure reveals a relatively hydrophilic surface unit, and a discrete hydrophobic region at the TMD (Figure 4C), as would be necessary for fusogenic function. Prediction of protein structure from candidate sequences show structural conservation with human syncytin-1 and an illustrative *percomORF* (Figure 4D). Like human syncytin-1, predicted structures from candidate sequences such as *S. canicula* and *Squalus* spp. show a canonical disulfide bond between the C-X-X-C and C-X_6_-CC motifs on the SU and TM units respectively (Figure 4E&F), linking the two. In the linear peptide sequence these cysteine residues are distant from each other (169 amino acids distant in synctin-1) and therefore their proximity to form a disulfide bond is a strong indicator that the 3D protein architecture and fusogenic function is functionally conserved. Although superficially similar, the predicted structure for *R. brachyura* shows no such bond, and the structure for *N. bancroftii* is ostensibly different. Overall, candidates from 9 species (6 sharks, 2 chimaeras, and 1 ray) were found to form canonical disulfide bonds in their 3D confirmation from the original 16 species with detected candidates (Figure 4F) and were carried forward for further investigation.

RNA-Seq datasets are sparse for chondrichthyans, with the majority of our species having poor transcriptomic coverage, however the small-spotted catshark *Scyliorhinus canicula* serves as a model organism for developmental studies [53, 54] and as such has a uniquely high coverage of tissue-specific transcriptomic datasets [55]. *S. canicula* was therefore used as a model system to examine whether syncytin-like Env candidates from chondrichthyans were expressed, and in which tissues. We searched for expression of our 21 *S. canicula* Env candidates in 245 RNA-Seq samples covering 16 tissue types. We found no expression of our candidates in any datasets derived from embryonic tissue. However, we detected their expression in nine diverse adult tissue types (Figure 5A; Supplementary File 3). Across tissues, the distributions were strongly zero-inflated, indicating that candidate transcription is intermittent rather than uniform, but that some tissue classes repeatedly harbour higher-expression loci. At the candidate level, expression was concentrated in a restricted subset of loci rather than being evenly distributed across all 21 candidates. Overall, 8 candidates exceeded a mean count of 10 in adult tissues, confirming that repeated moderate-to-high transcription is confined to a relatively small subset of loci (Figure 5B).

In intact retroviral proviruses, Env sequences are associated with nearby Gag and Pol genes necessary for viral replication [17]. However, disruption or absence of these contiguous genes around an intact Env sequence is evidence to support endogenization and exaptation into the vertebrate genome [17]. The majority (19/21) of *S. canicula* Env candidates had nearby reverse transcriptase support together with at least one additional Pol-associated domain, as would be predicted for a retroviral insertion event. At the domain level, reverse transcriptase, RNase H, and integrase/rve-like support were each present in all candidates, however Gag-like support was detected in only 2/21 loci. These data provide evidence that our candidates represent truly endogenized syncytin-like Env genes as opposed to intact proviruses (Figure 5C&D; Supplementary File 4). Phylogenetic clustering of the Pol genes associated with our *S. canicula* Env candidates with those from known retroviruses and syncytins provides evidence suggesting that candidates are descended from an ancestral Epsilonretrovirus (Figure 5E; Supplementary File 5).

If Env genes are detected in identical or similar chromosomal locations between closely related species, then this evidence supports inheritance from a common ancestral insertion event as opposed to repeated infection [56, 57]. Although the available chondrichthyan genomes we screened cover all major groups and nutritive strategies, they are at a relatively coarse taxonomic resolution between species. However, we detected Env candidates with canonical disulfide bonds in two closely related *Squalus* spp. (*S. suckleyi* and *S. acanthias*), which could be analysed for syntenic genome locations indicative of shared ancestry by whole-genome alignment. Ordering *S. suckleyi* scaffolds by their dominant *S. acanthias* chromosome assignment produced a coherent scaffold-to-chromosome synteny map, consistent with strong large-scale collinearity between the two *Squalus* spp. genomes (Figure 6). Projection of *S. suckleyi* Env candidates into the *S. acanthias* chromosomal space recovered cases of co-localisation where candidates overlapped (3), were nearby (2), were on the same chromosome (22), or were unresolved (5) (Figure 6A; Supplementary File 6). Of the five overlapping or nearby candidates, all were associated with present Pol genes but lacked Gag genes. Three of these loci were also retained within the filtered high-confidence alignment set and could therefore be visualised directly on the ordered synteny plot alongside their corresponding *S. acanthias* ORFs (Figure 6B). Taken together this evidence suggests true endogenization as analysed in *S. canicula*, as well as vertical transmission between *Squalus* species.

## Discussion

In this study, we mined chondrichthyan genomes, spanning the phylogenetic breadth and reproductive diversity of the entire clade, for the presence of syncytin-like retroviral Env sequences previously identified from mammals. Using strict bioinformatic criteria, we conservatively identified strong candidates for syncytin-like Env proteins in the genomes of nine non-placental chondrichthyan species, which contained the prerequisite domain architecture and 3D structure for fusogenic function. Using the catshark *S. canicula* as a model species, we demonstrated that many Env genes were expressed across diverse adult tissue types and showed clear signatures of endogenization into the host genome. In two closely related *Squalus* species, we showed that some Env sequences display high syntenic conservation, indicative of inheritance from a shared ancestral endogenization event prior to species divergence. Taken together, our clade-wide results show strong support that many of our candidate sequences represent fully endogenized and expressed syncytin-like Env proteins with fusogenic properties, expanding the known phylogenetic diversity of independent syncytin-like Env capture beyond mammalian placentation.

Most research into syncytin-like Env endogenization and function has centred on their role in mammalian placentation [22, 58]. By mediating cell-cell fusion of trophoblast cells into a syncytium, the independent capture of syncytins facilitates the uniquely intimate form of placentation found in many mammal species [17, 30], and knockouts of syncytin majorly disrupt placental formation [24, 59]. The centrality of retroviral capture on trophoblast fusion has focused research efforts into identifying novel functional syncytins in non-human mammal species, which have subsequently been identified in the placentas of mice [23], rabbits [60], squirrels [61], ruminants [56], carnivores [57], tenrecs [62], and marsupials [63]. The capture of a functional syncytin has also been shown to drive intimate placentation in the lizard *Mabuya* [64] and a syncytin-like Env gene is associated with hemochorial placentation in hyenas [30]. These detailed mechanistic studies have underscored the importance of independent syncytin capture in the evolution of intimate placentation. Our results inversely provide support for the association of syncytin-like Env capture with uniquely intimate placentation, as our pipeline detected no candidates in the genomes of placental sharks (four sharks in the Carcharhiniformes). As shark placentas are derived from the yolk sack and are epitheliochorial (not intimate with maternal circulation) [8, 65], the absence of syncytin-like Env candidates in shark placentation supports the hypothesis that syncytin capture is not associated with the independent evolution of less intimate placentation modes.

The capture of syncytin-like Env proteins outside of the context of mammalian placentation is less well understood. An ancient syncytin-like Env gene was detected in the genomes of teleost fish [31, 66], which has subsequently been shown to be a fusogenic protein (Percom-env) expressed in the brain [32]. Other endogenized teleost syncytin-like Env genes are expressed in gonadal tissue [32]. Syncytin-like Env candidates have also been identified in monotremes (egg-laying, non-placental mammals), which confer fusogenic properties and are highly expressed in tissues such as the kidneys [39, 67]. These important studies demonstrate a much wider role for endogenized syncytin-like Env proteins beyond mammalian placentation. Human syncytins have also been demonstrated to play a physiological role in non-placental contexts such as gamete fusion, osteoclast generation, and muscle fibre formation [25]. Our detection of syncytin-like Env candidates in non-placental sharks suggests that these proteins may play similar roles in chondrichthyan physiology. This is supported by our evidence that endogenized Env genes from *S. canicula* are expressed in adult brain, heart, and gonadal tissues, which corresponds to the previously demonstrated non-placental functions of teleost and human Env proteins [25, 32]. Our results that endogenized Env genes are likely vertically transmitted between *Squalus* species also implies a positive selective pressure with physiological function.

Our study emphasises phylogenetic breadth, and future studies will need to prioritise mechanistic depth of our detected candidates. Our work has identified a wide range of novel retroviral syncytin-like Env genes, many with demonstrable signatures of endogenization, expression, and syntenic conservation, however their physiological function is unclear for now. Future sequencing of additional chondrichthyan genomes at a higher taxonomic resolution, and the acquisition of corresponding tissue-specific transcriptomes, will help to better resolve the number of independent capture events and vertical transmission of syncytin-like Env genes between species, as well as the location of their expression within the animal. To fully examine whether our identified genes retain fusogenic function, antibodies must be raised against our candidates and *in vitro* transfection assays conducted, which are beyond the scope of our present bioinformatic study. Future *in vitro* and *in vivo* investigations into chondrichthyan Env function should begin with the catshark *S. canicula*, which as a model species in developmental biology is already supported by high-quality genomic resources and husbandry protocols, and in which our study identified fully endogenized and expressed syncytin-like Env candidates with domains indicative of fusogenic function. To conclude, our work, using a strict and robust bioinformatic pipeline, has identified Env genes with strong indicators of endogenization and fusogenic function and has laid a solid foundation for future research into independent syncytin capture beyond mammalian placentation.

## Data availability

All data used in this study are freely available as Supplementary Data alongside this manuscript.

