## Supplementary material for "Evidence for independent retroviral syncytin-like Env endogenization in non-placental chondrichthyans": _Supplementary_Data_Legends.txt

Supplementary File 1 - Aligned and trimmed FASTA file of known syncytin and syncytin-like query sequences used to identify chondrichthyan Env candidates.

Supplementary File 2 - Aligned and trimmed FASTA file of all 295 candidate syncytin-like Env sequences identified from our bioinformatic pipeline.

Supplementary File 3 - Expression counts for syncytin-like Env candidates in Scyliorhinus canicula tissue-specific RNA-Seq datasets. 

Supplementary File 4 - Results of the Gag/Pol domain analysis of Env candidates from Squalus spp. and S. canicula. 

Supplementary File 5 - Aligned and trimmed FASTA file of Pol genes from known retroviruses, syncytins, and S. canicula candidates used to build the tree in Fig. 5E

Supplementary File 6 - Results of the synteny analysis of detected syncytin-like Env candidates between Squalus acanthias and Squalus suckleyi. 
